# Single-nuclei transcriptomics of mammalian prion diseases identifies dynamic gene signatures shared between species

**DOI:** 10.1101/2022.09.13.507650

**Authors:** Athanasios Dimitriadis, Fuquan Zhang, Thomas Murphy, Thomas Trainer, Zane Jaunmuktane, Christian Schmidt, Tamsin Nazari, Jacqueline Linehan, Sebastian Brandner, John Collinge, Simon Mead, Emmanuelle Viré

**Affiliations:** MRC Prion Unit at University College London (UCL), Institute of Prion Diseases, UCL, London W1W 7FF, UK; Division of Neuropathology, the National Hospital for Neurology and Neurosurgery, University College London Hospitals NHS Foundation Trust; Department of Clinical and Movement Neurosciences and Queen Square Brain Bank for Neurological Disorders, Queen Square Institute of Neurology, University College London, London, UK; Department of Neurodegenerative disease, Queen Square Institute of Neurology, University College London, London, UK

## Abstract

Mammalian prion diseases are fatal and transmissible neurological conditions caused by the propagation of prions, self-replicating multimeric assemblies of misfolded forms of host cellular prion protein (PrP). The most common human form of the disease, sporadic Creutzfeldt-Jakob disease (sCJD), typically presents as a rapidly progressive dementia and has no effective treatments. Prion diseases are transmissible to laboratory rodents affording unprecedented opportunities to understand neurodegeneration in its evolving stages. Murine models are especially useful in prion research as they develop bona fide prion disease and recapitulate all biochemical and neuropathological hallmarks of human prion disease. Despite extensive studies investigating the changes in transcriptional profiles in prion diseases the mechanisms by which prion diseases induce cellular toxicity, including changes in gene expression profiles are yet to be fully characterized. This is at least in part because confounding effects related to brain cellular heterogeneity have not been resolved. Here, we took advantage of the recent developments in single-cell technologies and performed an unbiased whole-transcriptome single-nucleus transcriptomic analysis in prion disease.

## INTRODUCTION

Mammalian prion diseases are fatal and transmissible neurological conditions caused by the propagation of prions, self-replicating multimeric assemblies of misfolded forms of host cellular prion protein (PrP). The most common human form of the disease, sporadic Creutzfeldt-Jakob disease (sCJD), typically presents as a rapidly progressive dementia and has no effective treatments^1^. Prion diseases are transmissible to laboratory rodents affording unprecedented opportunities to understand neurodegeneration in its evolving stages. Murine models are especially useful in prion research as they develop *bona fide* prion disease and recapitulate all biochemical and neuropathological hallmarks of human prion disease^2^.

Despite extensive studies investigating the changes in transcriptional profiles in prion diseases^3-8^ the mechanisms by which prion diseases induce cellular toxicity, including changes in gene expression profiles are yet to be fully characterized. This is at least in part because confounding effects related to brain cellular heterogeneity have not been resolved. Here, we took advantage of the recent developments in single-cell technologies and performed an unbiased whole-transcriptome single-nucleus transcriptomic analysis in prion disease. In order to investigate the changes during the course of the disease, we used a well-characterised FVB mouse model intracerebrally inoculated with murine-adapted RML prions. Specifically, this model shows distinct phases of prion propagation, an exponential phase of increasing prion titre, followed by a levelling off which continues until clinical onset^9,10^.

## RESULTS

Mice were intracerebrally inoculated with RML or uninfected brain homogenate (N = 15 per group and time point), or sterile PBS (N = 5 per time point) and were culled at 5 time points corresponding to different levels of infectivity and stages of preclinical disease (20, 40, 80, 120 dpi and end-stage) (**Figure 1a**). In this model, prion titres increase exponentially for 20-80 days then reach a plateau level (**Extended Data Figure 1**). Spongiosis and abnormal PrP deposition were determined at the same time points and were exactly as expected from the Sandberg model^9^ (**Figure 1b**).

**Figure 1:**
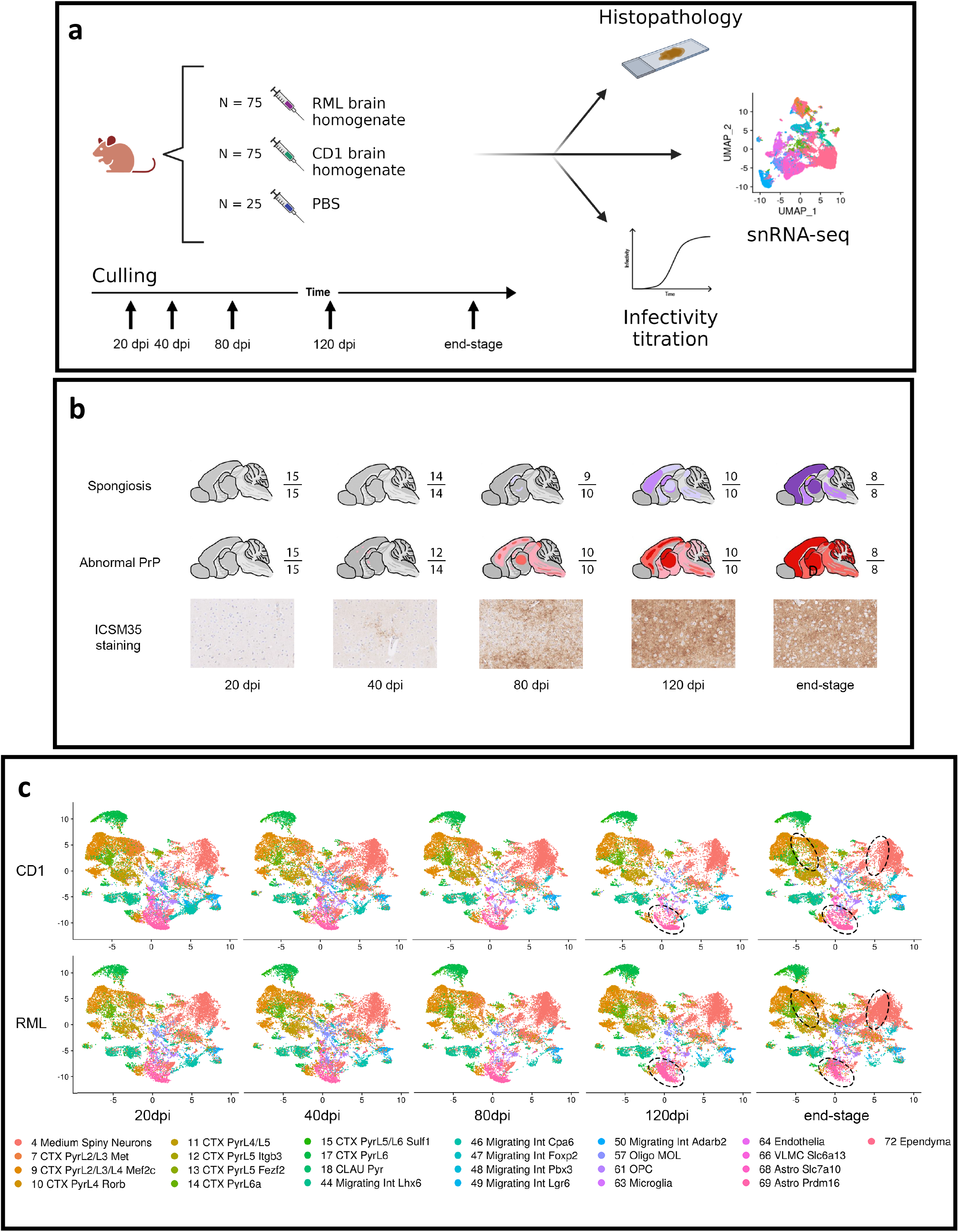
Longitudinal single-cell transcriptomics of the FVB RML-prion-infected mouse model. **a**, Experimental set up for the longitudinal single-cell transcriptomics study of prion disease in FVB mice. Mice were inoculated at day 0 with end-stage RML brain homogenate (N = 75), CD1 uninfected brain homogenate (N = 75), or PBS (N = 25). Samples were collected at time points corresponding to defined levels of prion infectivity (20 dpi, 40 dpi, 80 dpi, 120 dpi and end-stage). Brains were processed for single-nucleus transcriptomics, quantification of prion infectivity, and histopathological analyses. **b**, Formalin fixed brains from each time point underwent immunohistochemical analysis for abnormal PrP deposition and spongiosis. Schematic representations show the evolution of abnormal PrP deposition and spongiosis during disease progression. Top lane: overview of the distribution of vacuoles, where graded purple shades reflect the intensity of spongiosis. Mild spongiosis in the hippocampus and thalamus was evident in 9/10 animals at 80 dpi and became more pronounced in 10/10 animals as the disease progressed. The hippocampus showed mild neuronal loss (yellow) at clinical endpoint. Middle lane: overview of the distribution of abnormal prion protein deposits, where graded red shades reflect the intensity of prion protein deposits. Regarding the presence of residual inoculum in the early time points, a proportion of the brains (4/15) at time point 20 dpi, inoculated with RML, showed immunopositive material located at the fringe between hippocampus and corpus callosum. This material appeared as small, solid, and densely immunoreactive. Occasionally, there were processes, presumably from astrocytes of the hippocampus, which also showed weak immunolabelling. This finding suggests the existence of residual inoculum, and we interpreted the presence of immunoreactive material in astrocytes processes as an early prion uptake rather than *de novo* prion replication. A similar finding in 2/15 animals of the 40 dpi RML group was observed, however, there was additional fine granular immunopositive material, more in keeping with incipient, *de novo* production of the abnormal prion protein. In the subsequent time points (80 dpi, 120 dpi, and end-stage) no evidence of residual inoculum was identified. There was widespread *de novo* deposition of abnormal prion protein as expected in these timed culls. No immunoreactive material was seen in either of the 2 control groups (uninfected brain homogenate and PBS). Bottom lane: bright-field microscope images show the abnormal PrP deposition in the cortex of prion-infected mice using the ICSM35 antibody for staining. Abnormal PrP deposition became evident at 40 dpi, becoming more pronounced following disease progression. **c**, UMAP plots visualise the relationship between the 25 identified clusters in low-dimensional space across the 5 time points and suggest transcriptomic differences in neuronal and astrocytic populations at the last two time points. Most of the clusters comprise neuronal cells (cortical neurons, migrating interneurons and medium spiny neurons), while we also identified glia (mature oligodendrocytes, oligodendrocyte precursor cells, microglia, and astrocytes), endothelia, leptomeningeal cells, and ependymal cells. When the transcriptomes are split by experimental group, the first three time points (20dpi to 80dpi) do not show pronounced differences between the CD1 and RML groups. At 120dpi there was an observable shift of astrocytic populations (clusters 68 and 69; highlighted by ellipses). This difference became more pronounced at the end-stage where there was a more pronounced shift of the astrocytic populations (clusters 68 and 69; highlighted by ellipses), while there were also differences in the medium spiny neurons (cluster 4; highlighted by ellipses) and subpopulations of cortical neurons (clusters 7 and 9; highlighted by ellipses). Pyr: pyramidal; L2/L3/L4 etc.: layer 2,3,4 etc.; CTX: cortex/cortical; CLAU: claustrum; Int: interneurons; MOL: mature oligodendrocytes; OPC: oligodendrocyte precursor cells; VLMC: vascular and leptomeningeal cells; Astro: astrocytes.

We hand-dissected the frontal lobe of 105 samples (RML/CD1: N = 9, PBS: N = 3, at each time point) and processed for single-nucleus RNA sequencing with a SPLiT-seq protocol, adapted to allow safe work with prions in a Biological Safety Level 3 environment^11^. After extensive quality controls, clustering and annotation of the 210,710 high-quality transcriptomes (M = 1050 features per nucleus, SD = 565) we identified 25 subclusters of neurons, astrocytes, oligodendrocytes, oligodendrocyte precursor cells (OPCs), endothelia, microglia, ependymal and leptomeningeal cells based on location and the selective expression of transcription factors (**Figure 1c;** Error! Reference source not found.). Overall, dimensionality reduction plots suggested no visual transcriptomic shifts in RML-infected mice relative to CD1-inoculated mice up to and including 80dpi (**Figure 1c**). At 120dpi there was a visual hint of a shift in the transcriptomic profile of astrocytes (clusters 68 and 69; **Figure 1c**), which became more pronounced at the end-stage (**Figure 1c**) when differences were also observed in the transcriptomes of medium spiny neurons (cluster 4; **Figure 1c**) and subpopulations of cortical neurons (clusters 7 and 9; **Figure 1c**).

Next, we performed a differential gene expression analysis to identify specific transcriptomic alterations during disease progression. We identified a total of 928 differentially expressed genes (DEGs) across all five time points (log_2_-fold change > 0.25, Benjamini-Hochberg adjusted p-value < 0.05, Wilcoxon rank-sum test; Supplementary table 1). Importantly, the data suggested a selective transcriptomic response of individual cell clusters to disease, that could not be entirely accounted for by the cluster size and statistical power (Spearman correlation coefficient between cluster size and number of DEGs = 0.28). We found that at 120 dpi clusters 7 and 17 (cortical neurons) are early affected, and clusters 4 and 9 (medium spiny neurons and cortical neurons) seem quite unaffected relative to being the most common cell types by count. Similarly, at end-stage clusters 69 (Astro Prdm16), 72 (Ependyma) and 9 (Ctx PyrL2/L3/L4 Mef2c) are the most affected.

Longitudinal analysis in specific cell-types revealed a remarkable triphasic profile of gene expression changes as disease progresses. A first set of genes is affected as early as 20 dpi with clusters of cortical neurons, astrocytes and oligodendrocytes being mostly affected. Strikingly, this profile is not seen at 40 dpi nor at 80 dpi with only 6 and 5 genes respectively being affected. Then, a more substantial dysregulation in clusters is observed from the 120 dpi time point (173 genes in 12 clusters) and is amplified at end-stage with 683 genes dysregulated in 21 clusters. The highest number of identified DEGs were associated with astrocytes (cluster 69) and in a lesser degree mature oligodendrocytes (cluster 57) and oligodendrocyte precursor cells (cluster 61), especially in the last two time points. This reveals a pattern of oscillating transcriptomic perturbations commencing at 20 dpi, when infectivity was low, then subsiding at 40 and 80 dpi even though infectivity proceeded in an exponential fashion, before re-emerging during the infectivity plateau at 120 dpi and being amplified at the end-stage (**Figure 2a**).

**Figure 2:**
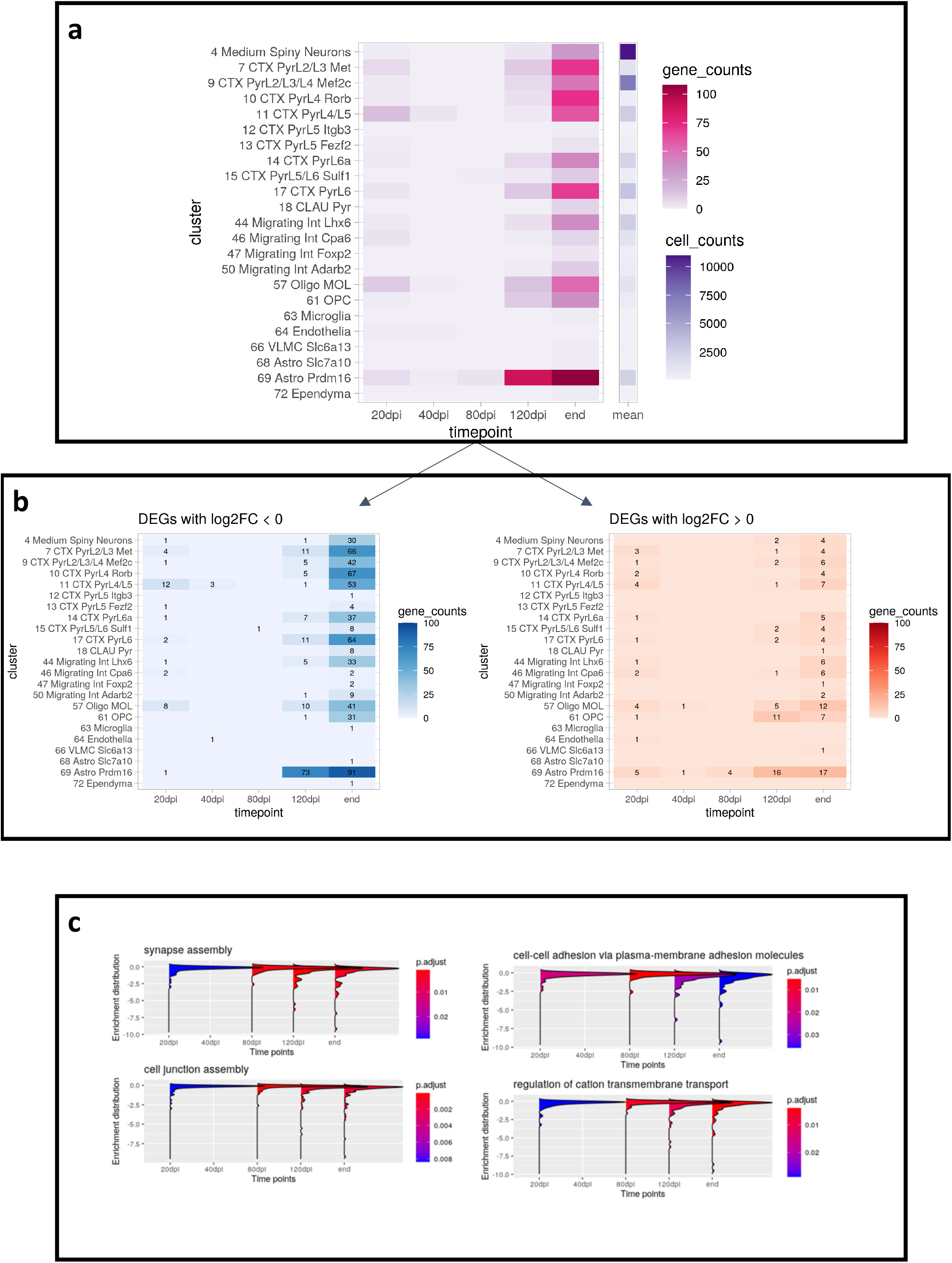
Single-cell transcriptomics reveals a tri-phasic gene expression landscape of prion disease’s progression. **a**, The heatmap shows the number of differentially expressed genes (DEGs) per cluster (y-axis) and time point (x-axis; in red), and the mean number of identified cells per cluster across all time points (in purple) when using Seurat’s non-parametric Wilcoxon rank sum test. The counts of differentially expressed genes and the counts of the cells in each cluster use different scales, which are denoted using two different colours. **b**, The two heatmaps separately represent the numbers of differentially expressed genes that decrease (left) or increase (right) in expression in response to disease across the 5 time points. The overall transcriptomic profile is shown to be primarily driven by down-regulated transcripts. **c**, The global suppression of transcription becomes progressively evident in multiple enriched Gene Ontology (GO) terms. The synapse and cell junction assembly, cell adhesion, and ion homeostasis GO gene sets showed gradual dysregulation that followed disease progression. These terms were shown to be dysregulated at the earliest time point, while no enrichment was observed at 40 dpi, before being enriched again at 80 dpi. The enrichment distributions followed a downward trend along with disease progression. Each plot represents a single GO term. The x-axis represents disease progression (left to right). The y-axis represents the distribution of the gene expression of genes associated with a specific GO term. Time points without plots indicate that the specific GO term was not enriched at that specific time point. The colour of each ridge plot represents the BH-adjusted p-value.

We found that at 20 dpi, 34 DEGs are downregulated and 26 are upregulated (**Figure 2b**). This balance shifts towards downregulation with 131 downregulated versus 42 upregulated genes at 120 dpi and 592 downregulated versus 91 upregulated genes at end stage (Supplementary table 1). Next we tested the differential expression signatures for enriched pathways and gene sets. We performed a gene set enrichment analysis (GSEA)^12^ using clusterProfiler which identified 4 pathways that showed an initial enrichment followed by progressive dysregulation, with gene expression diminishing as disease progresses (**Figure 2c**). This was the case for synapse assembly, cell function assembly; cell-cell adhesion via plasma-membrane adhesion molecules and regulation of cation transmembrane transport. Altogether, these findings highlight triphasic alteration to gene expression profiles in the course of prion disease progression, with most genes being downregulated as disease progresses, and brain-specific pathways being altered.

To further characterize the genes that are dysregulated in the course of prion disease, we dissected them into three sets with distinct transcriptional profiles (**Figure 3a**): 16 “early” genes showed dysregulation at 20dpi only; 393 “late” genes showed dysregulation from 120dpi and onwards; and 29 “resurgent” genes displayed an oscillating pattern of dysregulation whereby they were affected at 20 dpi and again at 120 dpi and onwards. Among the 16 early genes (**Figure 3b**), we found that *Nell1, Il1rapl2* and *Ppm1e* correlate with ageing of the brain^13^ and *PTPRD* is known to induce pre- and post-synaptic differentiation of neurons by mediating interaction with *IL1RAPL2* in human^14^.

**Figure 3:**
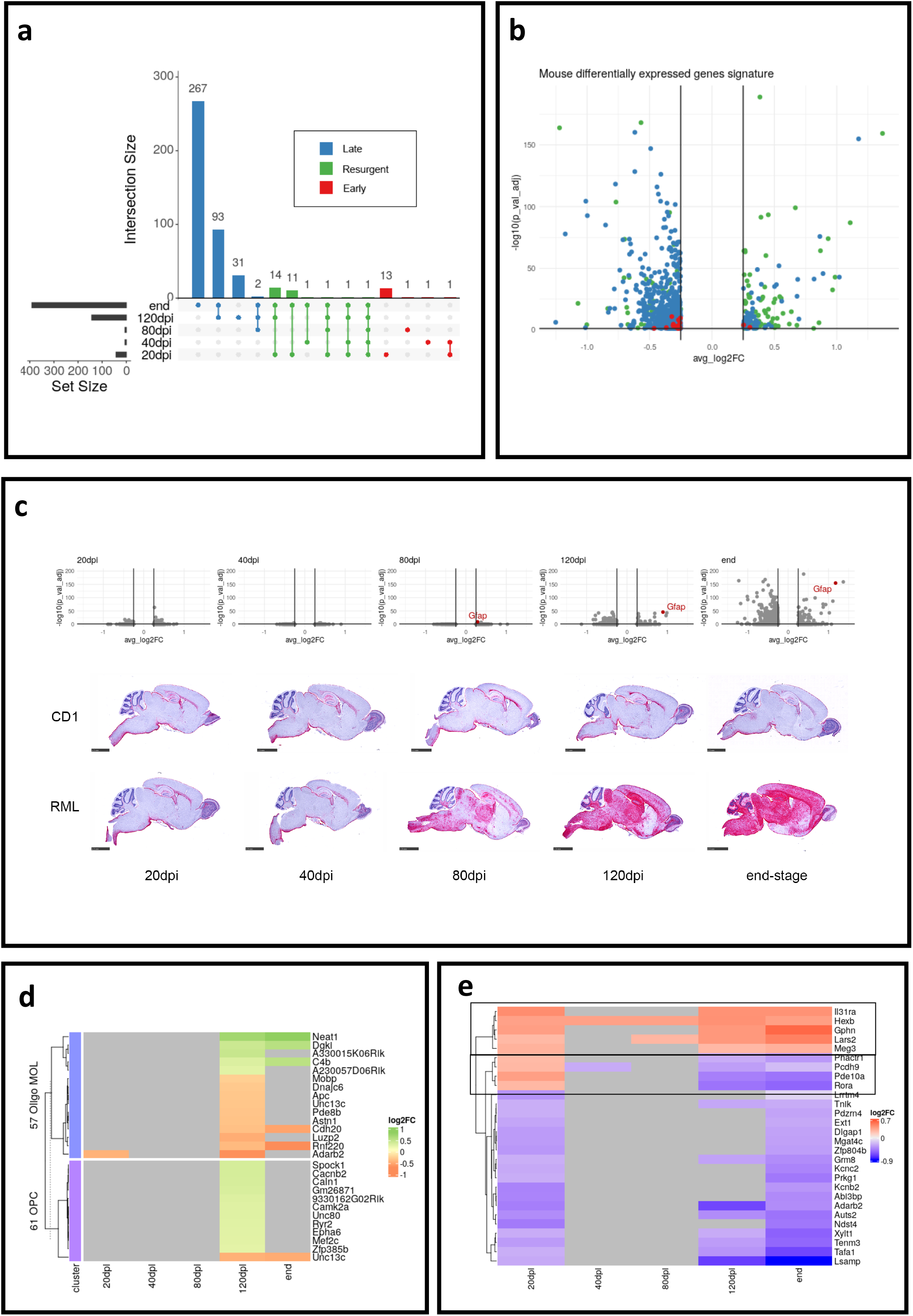
Characterization of early, late and resurgent genes in prion disease. **a**, UpSet plot shows the intersections between sets of DEGs across the 5 time points. **b**, Volcano plot of all DEGs at all 5 time points. Dysregulation of the “early” genes (red) is mild compared to the entirety of DEGs. “Resurgent” genes (green) and “late” genes (blue) are shown. The volcano plot includes all genes shown to be DE in any time point. The x-axis corresponds to the log-base-2-fold changes in gene expression. The y-axis corresponds to the minus log-base-10 of the BH-adjusted p-value. **c**, *Gfap* transcript increased from 80dpi. Top: volcano plots show *Gfap* in the snRNA-seq at each time point. Middle: Representative images of stained mouse brains using RNAscope showing increased *Gfap* expression in RML-inoculated mice as disease progresses. Each row corresponds to a different experimental group (top: CD1-inoculated mice, bottom: RML-inoculated mice), while each column corresponds to a different time point. Scale bars is length of 2.5 mm. Right: Quantification of *Gfap* RNAscope signals. Levels remain stable in control mice, except in the hippocampus, where *Gfap* levels decreased with ageing. A Shapiro-Wilks Normality Test was performed to ensure normality before calculating p-values using a two-way ANOVA. P-values were corrected using Sidak’s multiple comparisons test. Numbers on top of the bars represent the calculated p-values. N = 3 independent biological replicates per inoculum per time point. Points represent biological replicates. **d**, Heatmap of the DEGs associated with the clusters of mature oligodendrocytes (cluster 57) and OPC (cluster 61). **e**, Transcriptional profiles of the resurgent gene signature across the 5 time points. Heatmap of expression and clustering the genes (rows) based on their pattern of DE identified 3 groups: genes upregulated at 20dpi and in the later time points of the disease; genes upregulated at 20dpi, but downregulated at later time points; genes downregulated at 20 dpi and more strongly suppressed at the later time points.

In the resurgent genes, we found that *Gfap* upregulation started as early as 80 dpi. The *Gfap* signal at 80 dpi is consistent with staining for de novo deposition of abnormal prion protein with no immunoreactive material seen in the 2 control groups (normal brain homogenate and PBS) (**Figure 1b**). *Gfap* is considered to be a highly specific marker for astrocytes and is therefore often used to quantify their presence^15^. *Gfap* has previously been reported to be activated in prion disease^16^. To verify that this signal is not an artefact, we used RNAscope in situ hybridization to visualize *Gfap* in RML and CD1 inoculated animals at all time points (**Figure 3c**). We found astrocytic involvement initiating as early as 80 dpi, which mirrored the transcriptomic results with *Gfap* levels increasing from 80 dpi in the hippocampus, thalamus and the brainstem, while *Gfap* levels in the cortex (which was the tissue used for the transcriptomic study) remained lower until the 120 dpi time point. Astrocytes are the most affected cluster at 120dpi with 69 genes dysregulated. The second most abundant cluster is oligodendrocytes with 27 genes dysregulated in OPC and mature oligodendrocytes. Only 6 of these genes (*Neat1, Dgkl, C4b, Cdh20, Rnf220, Unc13c*) are also affected at end-stage suggesting an oligodendrocyte specific signature at 120 dpi. All but 1 OPC gene were upregulated (**Figure 3d**).

Finally, we explored the resurgent genes whose expression oscillates across the 5 time points. Hierarchical clustering of the genes reveals 3 distinct transcriptional patterns: upregulated at 20 dpi and upregulated as disease progresses; upregulated at 20dpi and downregulated as disease progresses; and downregulated at 20dpi and downregulated towards the end-stage (**Figure 3e**). Altogether our results identify a novel transcriptional signature of prion disease progression with a set of genes whose expression is affected at very early stage and then are dysregulated again at later stages of the disease.

Having used SPLiT-Seq in mice infected with RML prion, we next wanted to use it in human brain samples from patients with sporadic CJD. We carefully designed a case-control cohort of post-mortem and biopsy brain samples (Supplementary table 2). Human cortex samples were hand-dissected, white matter removed, and nuclei suspensions were prepared. Post-mortem sCJD brain samples, sCJD brain biopsies, control post-mortem brain samples and control brain biopsies were loaded on the same plate and processed in the same batch in order to reduce possible batch effects. Sequencing generated approximately 850 million reads in total, yielding a total of approximately 50,000 transcriptomes. Filtering and QC of the data eliminated up to 99% of the data (Supplementary table 3) rendering the interpretation of the analysis unfeasible. We therefore performed bulk RNA sequencing on the cohort of case-control post-mortem samples (5 sCJD vs 9 controls). We found 3,070 coding DEGs in sCJD versus matched controls samples (Supplementary table 4) (**Figure 4a**). Hierarchical clustering using the 50 most variable coding genes highlights the variability between samples (**Figure 4b**). A gene set enrichment analysis (GSEA) highlighted the noisiness of the dataset with the 5 top enrichment terms being unrelated to brain function (angiogenesis, embryonic development, regulation of angiogenesis, regulation of leukocyte activation and regulation of vasculature development) (**Figure 4c**). Despite the high level of variability between human samples, 96 DEGs identified in the single-cell longitudinal experiment in mice were found in the human dataset (**Figure 4d**), with 64.5% of those (62 genes) sharing direction of effect (downregulated in mice and downregulated in human). The bulk RNA sequencing in human post-mortem samples also identified early, resurgent and late genes (**Figure 4e**). Next we explored the gene ontology terms associated with the 96 genes shared between mouse and human datasets and identified brain specific signals with axonogenesis and positive regulation of cell projection organisation being the 2 most significant pathways identified (**Figure 4f**).

**Figure 4:**
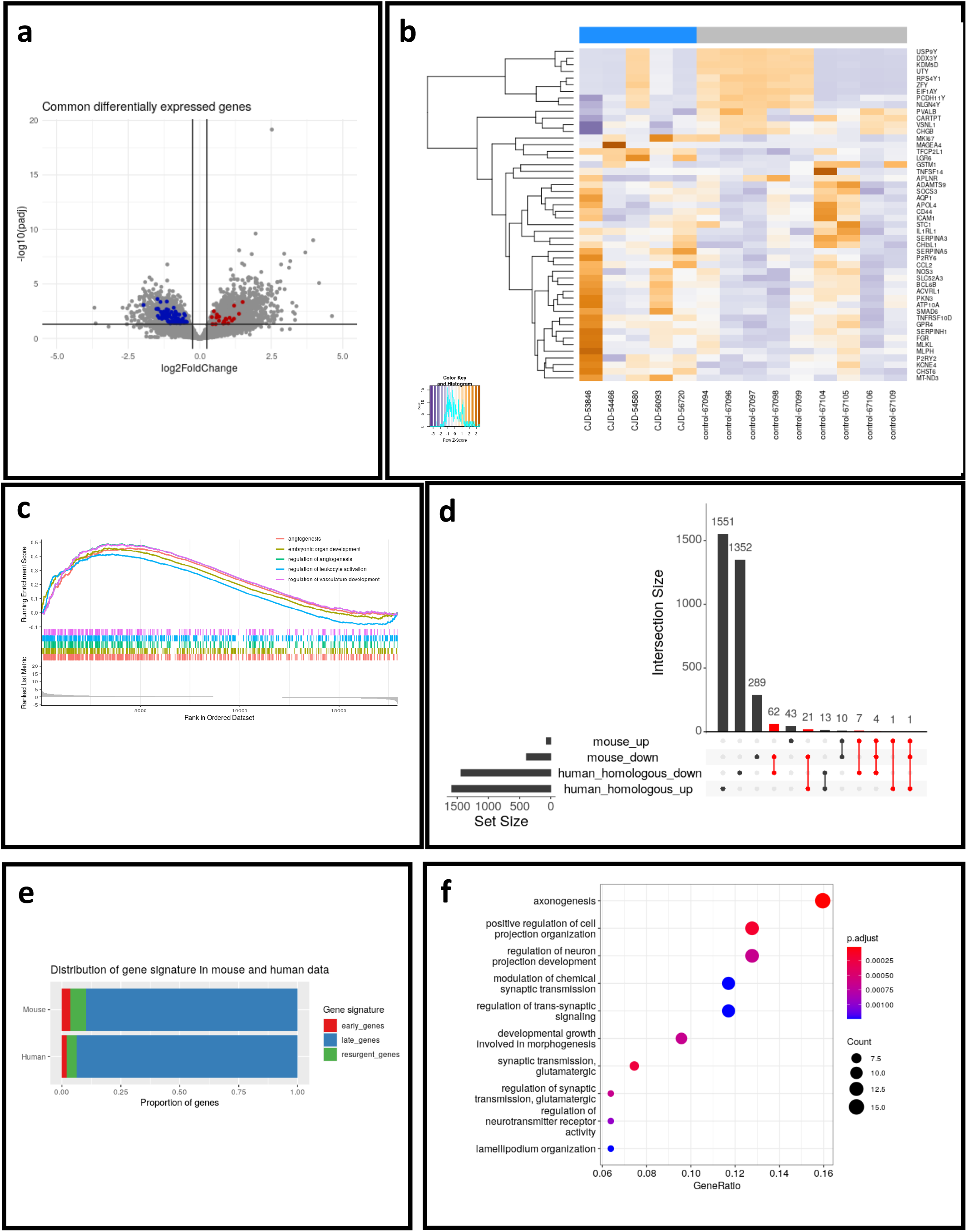
Early, late and resurgent genes are also found in sCJD brains. **a**, Volcano plot of all DEGs identified in the human dataset of sCJD (n=5) versus controls (n=9). Highlighted are the human coding genes with mouse homologues identified in the mouse single-cell study. Blue and red dots represent genes with decreased and increased expression, respectively. **b**, Hierarchical clustering of the top 50 most variable coding genes (rows) in sCJD (blue) and control brain samples (grey). **c**, Top 5 enriched terms as a result of a GSEA on DE genes of the human case-control study. **d**, UpSet plot of the intersections between mouse and human gene sets. **e**, Proportion of each of the 3 categories of genes (early, late, and resurgent genes) in human and mouse datasets. **f**, The common genes between the mouse and human datasets are associated with brain-specific Gene Ontology terms. The dotplot shows the results of a GO over-representation analysis on the intersection between the mouse and human datasets (mouse homologues).

## DISCUSSION

Here we performed unbiased transcriptomics analyses to gain insights into the gene signatures that characterize prion diseases in mammals. First, we used SPLiT-seq to study disease progression in a well-established mouse model of prion disease at a single-cell level. This led us to identify cell-type specific alterations in astrocytes, neurons and oligodendrocytes with distinct genes at early time points, later time points, or shared by early and late time points. This triphasic transcriptional landscape can be correlated with the time course of prion propagation^9,10^. At 20 dpi, SPLiT-seq detected changes in expression of 42 unique genes although no infectivity was detected using the automated scrapie-cell assay, nor was abnormal prion protein detected by histopathology. The changes in gene expression are most likely to represent a response of astrocytes, oligodendrocytes and neurons to the inoculation of toxic material associated with prion disease. Of note, all results discussed here are differential gene expression after comparison to inoculation with normal brain homogenate, therefore we can exclude the possibility that transcriptional changes resulted from any non-specific toxicity associated with inoculation of exogenous brain homogenate into the mice. Perhaps the most striking finding is the fact that transcriptional profiles remain almost unaffected at 40 and 80 dpi, the two time points where prion propagation grows exponentially implying a null transcriptional response to prion propagation itself. Yet, at later time points where signs of prion disease become visible histologically, and when prion propagation reaches a plateau phase, gene expression profiles are more strongly affected. This suggests that transcripts identified as differentially expressed in nuclei from RML inoculated animals correspond to a signature of prion-related disease. Our findings, where 28 transcripts are dysregulated both at very early and late time points reveals a subset of genes whose expression change may relate initially to toxicity in the inoculum, and later to that generated by the host disease process.

Unlike previous studies which investigated the translatome of major brain cell type categories^16^, our work considered the rich diversity of frontal cortex since the same number of nuclei were used as input from each sample at each time point. This enabled the assessment of disease-relevant cell types and disease progression specific transcriptomic signatures. Our findings complement the existing literature on transcriptomic alteration in prion diseases^3,5,8,16^.

We choose the SPLiT-seq method because it was compatible with the use of prion infected material. The caveat of that choice is that alternative methods would have provided more depth. As expected, gene lists could potentially be richer for each time point. Unlike in other neurodegenerative diseases, prion disease affects astrocytes regardless of their two canonical subtypes A1 (subtype being induced by neuroinflammation) or A2 (subtype referred to as neuroprotective astrocytes)^17,18^. Although astrocytic involvement in prion diseases has been demonstrated before^19^ here we reported changes to *Gfap* expression as early as 80 dpi in RML inoculated mice and visualized *Gfap* transcripts using an independent spatial technology. The increase in *Gfap* expression was seen in the entire brain using RNAscope with astrogliosis increasingly affecting all brain regions throughout disease progression.

Oligodendrocytes represent the second most affected cells in our dataset, yet, unlike astrocytes, which are known to replicate and accumulate prion independently of neurons^20^, oligodendrocytic contribution to cell-autonomous prion propagation is unlikely^21^. The alteration to oligodendrocytes is particularly evident at the 120 dpi time point with *Mobp* and at end-stage with *MBP* both downregulated. Bulk RNA sequencing from hand-dissected frontal cortex from sCJD cases and controls revealed an overlap in transcriptomic signatures between mice and human prion diseases, with 96 genes whose expression is dysregulated in both species. This suggest that despite fundamental differences in the technique and disease model, brain cells share similar molecular pathways of response to prion diseases. It is well established that single-nuclei technologies do not allow to study the contribution of microglia^22^, therefore we are unable to establish whether microglia is affected here.

In summary, our work shows the cell-type and disease stage-related gene expression changes during the time course of mammalian prion disease. We describe what appears to be an initial transient response to prion-disease brain inoculation, followed by a silent phase of unaltered gene expression during exponential prion propagation, and finally, a partly resurgent signature as neurodegeneration takes hold, led by changes in glial cell-types and specific neuronal clusters. Shared gene signatures between mice and human prion diseases highlight the potential of single-cell studies to better characterize the molecular mechanisms at the core of prion diseases.

## Acknowledgments

We are grateful to Malin Sandberg, Stephanie Canning, Dr Azadeh Khalili-Shirazi for providing technical assistance. We are very grateful to patients and volunteers who have made this study possible by donating samples and their time to research; Holger Hummerich for helpful discussions; Tony Brooks, Paola Niola, and Mark Kristiansen from UCL Genomics for technical assistance with sequencing; Tracy Campbell and Penny Norsworthy for help with metadata acquisition; Richard Newton for help with preparation of figures. We thank Prof. Goncalo Castelo-Branco for their comments and suggestions during the data analysis steps; Richard Newton for preparation of figures; Simran Singh for technical assistance with slides scanning. This work was funded by Alzheimer’s Research UK (ARUK-PPG2020A-030) to E.V. A.D. is supported by the Onassis Foundation - Scholarship ID: F ZQ 022-1/2020-2021. ZJ and SB are supported by the Department of Health’s NIHR Biomedical Research Centre’s funding scheme to UCLH. S.M. and J.C. are NIHR Senior Investigators.

## Methods

### Mice

All animal work was performed under approval and license granted by the UK Home Office (Animals (Scientific Procedures) Act 1986), which conformed to UCL institutional and Animal Research: Reporting of In Vivo Experiments (ARRIVE) guidelines. Experimental design adhered to the principles of the 3Rs - Replacement, Reduction and Refinement. 4–6-week-old female FVB inbred mice (FVB/NHan Hsd; Envigo) were intracerebrally inoculated around 6-8 weeks old. Inoculations on anaesthetised mice in the right parietal lobe with 30 uL of one of the following preparations: 1% RML prion-infected brain homogenate prepared from 10% RML stock; 1% uninfected CD1 brain homogenate prepared from 10% uninfected CD1 stock; 1x sterile DPBS (Gibco; 14190-086).

Mice were monitored daily for neurological signs of the disease and were culled by CO_2_ exposure either on schedule for the 20, 40, 80 and 120 dpi time points, or at scrapie sickness confirmation for the end-stage. Early signs include erect ears, rigid tail, piloerection, and ungroomed appearance, slight hunched posture, and clasping of hind limbs when lifted, while scrapie was confirmed when signs of ataxia, generalized tremor, loss of righting reflex, or limb paralysis was observed. When culling, the brain was removed and the left hemisphere was stored in 10% formal saline for further histopathological analysis, while the right hemisphere was snap frozen and stored at -80°C until further processing.

### Immunohistochemistry for prion-related neuropathology

Immunohistochemistry was performed as previously described with modifications^23^. Briefly, mouse brains were fixed in 10% buffered formal saline and paraffin wax embedded. Serial sections of 5 um were taken and deparaffinised. The sections were then processed to investigate PrP deposition on a Ventana Discovery XT automated IHC staining machine (Roche Tissue Diagnostics) using protocols developed on a Ventana Benchmark staining machine^24^. Sections were treated with cell conditioning solution (Discovery CC1; Roche Tissue Diagnostics) at 95°C for 60 minutes or with a medium concentration of protease (Protease 1; Roche Tissue Diagnostics) for 4 minutes. For PrP deposition anti-PrP monoclonal antibodies ICSM35^25,26^ were used in conjunction with biotinylated polyclonal rabbit anti-mouse immunoglobulin secondary antibodies (Dako; Agilent) and Ventana proprietary detection reagents utilizing 3,3′-diaminobenzidine tetrahydrochloride as the chromogen (DAB Map Detection Kit; Roche Tissue Diagnostics). For haematoxylin and eosin (H&E) staining conventional methods on a Gemini AS Automated Slide Stainer (Thermo Fisher Scientific) were used. Positive controls for the staining technique were used throughout. All slides were digitally scanned on a Hamamatsu NanoZoomer 360 instrument, and images were captured from the NDP.serve3 software (NanoZoomer Digital Pathology) and composed with Adobe Photoshop.

### Preparation of brain homogenates

2 mL screw cap tubes with conical bottom (Alpha Laboratories; CP5932) were filled with ribolysing beads (Fisher Scientific; 15515809) to cover the bevelled bottom of the tube and weighted. One frozen right mouse brain hemisphere was transferred in each tube and tubes were weighted again to calculate the mass of the brain. Appropriate volume of PBS was then added (Gibco; 14190-086) to prepare a 20% w/v homogenate (x4 the brain mass, assuming that brain tissue density is close to 1). The tubes were then tightly screwed, and tissue was homogenised in a Precellys Evolution homogeniser (Bertin Instruments; P000062-PEVO0-A) operated at 6500 rpm for 45 seconds. The tubes were then left at 4°C for 1 h to reduce frothing. 500 uL of homogenate was pipetted out and transferred in a new tube, where it was diluted to 10% w/v using PBS. Homogenates were stored at -80°C.

### Scrapie Cell Assay

The scrapie cell assay was performed as previously described, in an automated manner^27^. Briefly, the cell lines were seeded at 1.8*10^4^ cells / well in a 96-well plate, 24 hours prior to infection with 10% w/v RML brain homogenate at the following dilutions: 3*10^−6^, 10^−6^, 3*10^−7^, 10^−7^, 3*10^−8^, 1*10^−8^. The cells were then split using an automated liquid handling robot (Beckman Coulter; Biomek FX) every three to four days and assays after the third and fourth splits. 25,000 cells were plated on ELISpot IP Filter Plates (PVDF membrane, 0.45 um, Merck; MSIPN4550) and fixed at 50°C for 1 h before treatment with 1 ug/ml proteinase K (Roche; 3115828001) in lysis buffer (50 mM Tris HCl pH 8, 150 mM NaCl, 0.5% w/v sodium deoxycholate, 0.5% v/v Triton X-100) at 40°C for 1 h. Plates were washed and treated with 3M guanidine thiocyanate (Melford; G54000) for decontamination and antigen retrieval before blocking with SuperBlock blocking buffer (Thermo; 37545). Staining was performed using anti-PrP antibody (clone ICSM18; D-Gen Ltd) followed by detection with alkaline-phosphatase-linked anti-IgG1 antiserum (Southern Biotech; 1070-04). Spots were visualised with alkaline phosphatase conjugate substrate (Bio-Rad; 170-6432) and PK-resistant infected cells were counted using the Bioreader 5000-Eβ (BioSys Karben, Germany).

### Preparation of single-nuclei suspensions from frozen mouse brain

Flash-frozen mouse brain was left to partially thaw. The olfactory bulb removed, and a small slice of the frontal cortex was cut and transferred to a glass 2 mL dounce tissue homogenizer on ice. All following steps were carried out on ice and using ice-cold solutions. All centrifugation steps were performed at 500 g for 5 minutes at 4°C using a pre-chilled centrifuge, unless otherwise specified. 2 mL of Nuclei EZ prep was added, and tissue was homogenised using 20 strokes of the loose and 20 strokes of the tight pestle. The suspension was transferred to a 15 mL tube and 2 mL of Nuclei EZ prep was added. The suspension was incubated for 5 minutes and then centrifuged. The supernatant was discarded, 4 mL of Nuclei EZ prep was added, and the pellet was resuspended using a P1000 pipette. The suspension was incubated for 5 minutes and then centrifuged. The supernatant was discarded, and the pellet was resuspended in 4 mL Nuclei Suspension Buffer (NSB; 1x PBS, 0.01% BSA (Cambridge Bioscience; 227-10210) and 0.1% NxGen RNAse inhibitor (Lucigen; 30281-2)). The suspension was centrifuged, the supernatant discarded, and the pellet resuspended in 1 mL NSB. The suspension was filtered through a 35 um filter (Fisher Scientific; 10585801), and stored on ice.

For nuclei fixation, 3 mL of 1.33% formalin solution were added to the 1 mL of nuclei suspension. The suspension was incubated on ice for 10 minutes. 169 uL of 5% Triton-X was added to the fixed nuclei, the solution was mixed by pipetting and then incubated on ice for 3 minutes. Nuclei were centrifuged at 500g for 5 minutes at 4°C, supernatant was discarded, and the pellet was resuspended in 500 uL cold NSB. 500 uL of cold 100 mM Tris and 20 uL of 5% Triton-X were added. Nuclei were centrifuged again under the same conditions; supernatant was discarded, and the pellet was resuspended in 400 uL of cold 0.5X PBS. Nuclei were filtered through a 35 um filter, counted using a Neubauer-Improved haemocytometer, and diluted to a final concentration of 2000 nuclei/uL using cold 0.5x PBS supplemented with 0.2 units/uL SUPERase In RNAse inhibitor. The nuclei were stored frozen at -80°C until library preparation.

### Nuclei extraction from frozen post-mortem human brain and human brain biopsies

For the post-mortem samples, each human brain was left to partially thaw and removed from the storage cassette. A small slice of the superior frontal gyrus (approximately 50-100 mg) was cut and transferred to a glass 2 mL dounce tissue homogenizer on ice. For the human biopsies, no structure was visible, and a small slice (approximately 50-100 mg) was transferred to a glass 2 mL dounce tissue homogeniser. All following steps were carried out on the ice and using ice-cold solutions. All centrifugation steps were performed at 500 g for 5 minutes at 4°C using a pre-chilled centrifuge unless otherwise specified. 1.5 mL of Nuclei EZ prep was added, and tissue was homogenised using 20 strokes of the loose and 20 strokes of the tight pestle. The suspension was transferred to a 2 mL tube and 0.5 mL of Nuclei EZ prep was added. The suspension was incubated for 5 minutes and then centrifuged. The supernatant was discarded, 2 mL of Nuclei EZ prep were added, and the pellet was resuspended using a P1000 pipette. The suspension was incubated for 5 minutes and then centrifuged. The supernatant was discarded, 0.5 mL of wash buffer (1x PBS, 1% BSA (15260037; Gibco), 0.2 u/μL SUPERase In (AM2694; Invitrogen)) were added without resuspending, and the sample was incubated for 5 min to allow buffer interchange. Then 1.5 mL of wash buffer was added, and the sample was resuspended. The suspension was centrifuged, the supernatant was discarded, the pellet was resuspended in 500 μL wash buffer, and 0.5 mL of 50% OptiPrep Density Gradient Medium solution (D1556; Sigma-Aldrich) was added. The suspension was transferred on top of a 1 mL 29% OptiPrep cushion solution in a new tube and centrifuged at 10,000g for 30 min at 4°C. The supernatant was discarded, and nuclei were resuspended in 750 μL Parse Nuclei Buffer (from the Parse Evercode WT kit) + 0.75% Bovine Albumin Fraction V (15260037; Gibco). Preparation then proceeded following step 7 of the Parse nuclei fixation protocol (page 15 of the protocol).

The following steps were performed using the Parse Evercode nuclei fixation kit (Parse Biosciences) according to the manufacturer’s instructions.

Fixed nuclei were counted using a Neubauer-Improved haemocytometer and diluted to variable concentrations calculated using the sample loading table provided using cold nuclei suspension buffer provided with the Parse Evercode nuclei fixation kit (Parse Biosciences). The nuclei were stored frozen at -80°C until library preparation.

### Single-nucleus library preparation using SPLiT-seq

Sequencing libraries from the mouse brain samples were prepared using the SPLiT-seq protocol (version 3)^11^. A summary of the methodology and optimisations is given below.

For the reverse transcription, 4 uL the first 24 wells of Stock plate 1 were transferred to a new PCR plate on ice. 8 uL of RT mix (per reaction: 4 uL Maxima 5x RT buffer, 0.124 uL NxGen RNAse inhibitor, 0.25 uL SUPERase In, 1 uL 10 mM Takara dNTPs, 2 uL Maxima H minus enzyme, 0.625 uL H_2_O) was added to each well and then 8 uL of fixed nuclei suspension. The plate was placed in a thermocycler and PCR was carried out as per protocol instructions. All wells were then pooled together, Triton-X was added to a final concentration of 0.1% and suspension was centrifuged for 3 minutes at 500g. The supernatant was discarded, and nuclei were resuspended in 2 mL 1x NEBuffer 3.1 (New England Biolabs; B7203S) + 20uL NxGen RNase Inhibitor.

For ligation round 1, the ligation mix (1337.5 uL water, 500 uL 10x T4 ligase buffer (included with ligase enzyme), 100 uL T4 DNA Ligase (New England Biolabs; M0202L), 100 uL BSA 10 mg/mL, 12.5 SUPERase In, 40 uL NxGen RNAse inhibitor) was added to the nuclei suspension and into a basin. 40 uL of the suspension were pipetted in each well of the Ligation round 1 barcode plate and the plate was incubated in a plate shaker at 37°C for 30 minutes and 300 rpm rotation. Then 10 uL of the Ligation Round 1 blocking solution (316.8 uL 100 uM BC_0216, 300 uL 10x Ligase buffer, 583.2 uL water) were added to each well and the plate was incubated again under the same conditions.

For ligation round 2, the nuclei suspensions were pooled in a 15 mL Falcon and passed through a 40 um strainer to another Falcon. 100 uL T4 DNA ligase was added and the mix was transferred to a basin. 50 uL of the mix was added to each well of the Ligation Round 2 barcode plate. The plate was incubated as previously. Then 20 uL of the Ligation Round 2 blocking solution (369 uL 100 uM BC_0066, 800 uL 0.5 M EDTA, 2031 uL water) was added to each well. The wells were pooled in a 15 mL Falcon and passed through a 40 um strainer into another Falcon.

70 uL 10% Triton-X was added to the mix and nuclei were centrifuged for 5 minutes at 1000g. The supernatant was aspirated, and nuclei were washed with 4 mL wash buffer (4 mL 1x PBS, 40 uL 10% Triton-X, 10 uL SUPERase In) and centrifuged again under the same conditions. Supernatant was aspirated and nuclei were resuspended in 100 uL 1x PBS + 2 uL SUPERase In. Nuclei were counted using a Neubauer-Improved haemocytometer and a desired number of them was aliquoted in each 1.5 mL Eppendorf tube. PBS was used to fill each tube up to 50 uL. Each tube is referred to as a “sublibrary”.

For nuclei lysis, 50 uL 2x lysis buffer (final concentrations: 20 mM Tris pH 8, 400 mM NaCl, 100 mM EDTA pH 8, 4.4% SDS (Thermo Fisher Scientific; AM9822)) were added to each tube followed by 10 uL 20 mg/mL Proteinase K. The mix was incubated at 55°C for 2 hours with shaking at 300 rpm. Lysates were frozen at -80°C and processed the following day.

5 uL 100 uM AEBSF (Abcam; ab141403) was used to stop proteinase reaction. Dynabeads MyOne Streptavidin C1 (Thermo Fisher Scientific; 65002) were washed and used to bind the barcoded transcripts as per protocol. Beads were resuspended in 200 uL Template Switch mix (88 uL water, 44 uL 5x Maxima buffer, 44 uL 20% Ficoll PM-400, 22 uL 10 mM Takara dNTPs, 5.5 uL NxGen RNAse inhibitor, 5.5 uL 100 μM Template Switch Oligo, 11 uL Maxima H minus enzyme) and incubated at a rotating incubator at room temperature for 30 minutes and at 42°C for 1.5 hours.

The sample was then washed, resuspended in PCR mix (121 uL 2x Kapa Hifi Master Mix, 9.68 uL 10 uM BC_0108, 9.68 uL 10 uM BC_0062, water up to 242 uL) and split equally in 4 different wells of a PCR plate. The following PCR programme was then used: 3 min at 95°C, then 20 s at 98°C, 45 s at 65°C, 3 min at 72°C for a total of 5 cycles, and then hold at 4°C. Reactions were combined in a single tube, cleaned with 0.6x AMPure XP beads as per manufacturer’s instructions, eluted in 20 uL water, mixed with 180 uL of the same PCR mix and split in 4 wells (50 uL per well). 2.5 uL EvaGreen (Biotium; #31000) was added to each well and amplification continued in a QuantStudio 7 Flex qPCR machine (Thermo Fisher Scientific) until signal plateaued out of exponential amplification using the following programme: 3 min at 95°C, then 20 s at 98°C, 20 s at 67°C, 3 min at 72°C until signal plateaus out of exponential amplification, then 5 min at 72°C, hold at 4°C. Reactions were combined in a single tube, cleaned with 0.6x AMPure XP beads as per manufacturer’s instructions, eluted in 10 uL water and analysed at TapeStation 2200 using a gDNA tape. 600 pg of each sample was used for tagmentation using the Nextera XT sample prep kit using custom primers one of BC_0076-BC_0083 and BC_0118, according to manufacturer’s instructions. Resulting libraries were analysed on a TapeStation 2200 using a high sensitivity D1000 tape or a high sensitivity D5000 tape.

### Evercode Whole Transcriptome library preparation

Evercode WT (whole transcriptome) is the proprietary and optimised protocol that evolved from SPLiT-seq. The methodology is closely related to that of SPLiT-seq, with a few differences. The Parse Evercode Whole Transcriptome kit (Parse Biosciences) contains all consumables and a detailed protocol that includes all steps from nuclei fixation up to sequencing, including catalogue numbers of reagents. All steps were carried out according to manufacturer’s instructions following the official protocol.

Suspension preparation and split-pool barcoding were carried out in a BSL-3 laboratory. To move the sample to a BSL 2 laboratory, the following prion decontamination procedure was followed: at the end of the Parse Evercode WT user manual chapter 3.4, the resulting solution was incubated with 3 volumes of TRI-reagent at room temperature for 2 hours (R2050-1-50; Zymo Research). The mix was then transferred out of the BSL-3 facilities and nucleic acids were purified using the Direct-zol DNA/RNA miniprep kit (R2080; Zymo Research). The kit columns were substituted with Zymo-Spin IC Columns (C1004-50; Zymo Research) so that smaller elution volumes could be used, following the advice of Zymo customer support. 11 μL of the RNA fraction and 10 μL of the DNA fraction were eluted in the same tube. The resulting solution of 21 μL was used for the PCRs starting at section 3.5 of the Parse Evercode WT user manual.

### Next-generation sequencing

Resulting libraries were sequenced on an Illumina NextSeq 500 using a NextSeq 150 cycle Mid Output kit (Illumina; 20024904) according to manufacturer’s instructions. The settings used were: paired-end reads, read 1 length: 66 nt, read 2 length: 94 nt, Index 1 length: 6 nt.

### Single-cell data analysis

Fastq files were subjected to quality control using a dockerised version of FastQC^28^ pulled from the repository biocontainers/fastqc:v0.11.9_cv8. The generated reports were manually examined for sequencing quality.

For SPLiT-seq, the fastq files were aligned to the human transcript using STAR aligner^29^. The resulting sam files were converted to a binary format and were processed by the SPLiT-seq bioinformatics open-source tools (https://github.com/yjzhang/split-seq-pipeline) to generate a count matrix. Transcriptome GRCm39 (mm39) was used for aligning the mouse data. The count matrix generated was used as an input for the subsequent analyses.

For Evercode, the fastq files were processed using the Parse Biosciences pipeline v0.9.6 to generate the counts matrix. Its function and processes are similar to the SPLiT-seq open-source tools, and the final output is a count matrix that is used for further analysis.

For the analysis of mouse and human data, a pipeline based on the Seurat v4 R package was employed, following the official vignettes and recommendations^30-32^. A summary of the methodology is provided below, while the analysis scripts will be available online at our GitHub repository: https://github.com/athanadd/.

The count matrices generated were first used to create Seurat objects and relevant metadata regarding was added. Then Ensembl IDs were converted to gene symbols using EnsDb version 104.

### Single-cell data quality control

For quality control, the cells were filtered on the number of features to exclude low quality cells and possible duplicates with a low threshold of 250 and a high of 2500. The percentage of mitochondrial genes was calculated and cells with more than 1% mitochondrial genes were discarded. A cell cycling score for the S and G2/M phases was assigned using known cell cycling genes (MCM5, PCNA, TYMS, FEN1, MCM7, MCM4, RRM1, UNG, GINS2, MCM6, CDCA7, DTL, PRIM1, UHRF1, CENPU, HELLS, RFC2, POLR1B, NASP, RAD51AP1, GMNN, WDR76, SLBP, CCNE2, UBR7, POLD3, MSH2, ATAD2, RAD51, RRM2, CDC45, CDC6, EXO1, TIPIN, DSCC1, BLM, CASP8AP2, USP1, CLSPN, POLA1, CHAF1B, MRPL36, E2F8 as gene markers for the S phase, and HMGB2, CDK1, NUSAP1, UBE2C, BIRC5, TPX2, TOP2A, NDC80, CKS2, NUF2, MKI67, CENPF, TACC3, PIMREG, SMC4, CCNB2, CKAP2L, CKAP2, AURKB, BUB1, KIF11, ANP32E, TUBB4B, GTSE1, KIF20B, HJURP, CDCA3, JPT1, CDC20, TTK, CDC25C, KIF2C, RANGAP1, NCAPD2, DLGAP5, CDCA2, CDCA8, ECT2, KIF23, HMMR, AURKA, PSRC1, ANLN, LBR, CKAP5, CENPE, CTCF, NEK2, G2E3, GAS2L3, CBX5, CENPA as gene markers for the G2/M phase), and cell separation based on cell cycle was assessed by examining the PCA plots. Cell cycle was not regressed out.

### Single-cell data normalisation

The Seurat object was then split by experimental group (CD1, RML, PBS for the mouse experiment, sCJD and Control for the human experiment) and individual objects were normalised using SCTransform. The objects were then combined in one integrated object by first selecting the integration features and finding integration anchors. The integrated object was then annotated using label transfer from an annotated reference dataset.

### Single-cell data annotation/label transfer

For the annotation of the mouse data, the first step was to pre-process the reference data to be used for label transfer and cluster annotation. We used the published SPLiT-seq mouse data as a reference as it is well annotated and perfectly matches the sequencing methodology. Postnatal day 2 and 11 data obtained from mouse brain was downloaded from GEO (Sample GSM3017261) and filtered to include only anatomical regions that are found in the frontal lobe. We then used Seurat to normalise the datasets using SCTransform, select the integration features using the top 3000 variable features and prepare the integration anchors. The dataset was integrated, and a Principal Component Analysis was used to identify the first 50 PCs. The FindTransferAnchors and TransferData functions were used to identify data transfer anchors and transfer cell type metadata from the annotated reference to our datasets. The predicted cluster scores and mapping scores were visualised by generating histograms and clusters consisted of less than 100 cells were removed. The data was normalised again using SCTransform and PCs and UMAP coordinates were calculated.

The success of the reference data label transfer was assessed by plotting the expression of known marker genes in each cell type (Aqp4, Slc1a2, Plpp3, Gja1 for astrocytes; Mbp, Plp1 for oligodendrocytes; Vcan, Mbp, Pdgfra for oligodendrocyte precursor cells; Rgs5, Flt1, Ly6c1, Pltp for endothelial/smooth muscle cells; Dock2, Dock8, Csf1r, P2ry12 for microglia/macrophages; Dnah11 for ependymal cells; Gria1, Snhg11 for neurons), and statistics such as the number of cells, the mean of features and the mean of counts in each cluster were calculated.

### Single-cell data differential gene expression using Seurat

Differential gene expression between the same two clusters across different conditions was performed using Seurat’s FindMarkers function. The statistical test used was the non-parametric Wilcoxon rank sum test and the adjusted p-value was based on Bonferroni correction using all features in the dataset. The differentially expressed genes were then filtered, keeping the ones that had an adjusted p-value of less than 0.05.

For the mouse dataset, to generate the differentially expressed gene lists of the RML versus CD1 groups, initially a comparison between CD1 and PBS groups was done to identify genes that are shown to be dysregulated but could be due to technical noise or relevant to the inoculation with a brain homogenate and not prion specific. From the list of those genes, we selected genes that were identified in multiple clusters (more than 5) and excluded them from the RML vs CD1 comparison to reduce technical noise. This resulted in 7 excluded genes which were: Calm1, Cdk8, Cmss1, Malat1, mt-Rnr1, mt-Rnr2, and Rn18s.

For the human dataset, a full comparison of sCJD vs controls was done, without the exclusion of any genes.

### Single-cell data Gene Ontology over-representation analysis (ORA) and Gene Set Enrichment Analysis (GSEA)

We used clusterProfiler in R for both ORA and GSEA^33^. For the ORA we used the enrichGO function using all the available features in the dataset as the gene universe and the filtered differentially expressed genes as the query genes. The adjusted p-values were calculated using the Benjamini-Hochberg method. GO terms that were supported by less than 3 genes were filtered out. The AH92582 annotation database was used for this analysis.

For the GSEA we used the gseGO function with the same organism databases and setting the minimal size of each gene set to 10, the maximal size of genes annotated for testing to 500, and the p-value cut-off to 0.05. The adjusted p-values were calculated using the Benjamini-Hochberg method.

### RNAscope

mRNA was detected as red punctae in coronal FFPE mouse brain sections counter-stained with haematoxylin using RNAscope 2.5 VS target probes (Advanced Cellular Diagnostics; ACD) against each transcript (*Prnp*, Cat No. 476619; *Gfap*, Cat No. 313249; *C3*, Cat No. 417849) and an RNAscope VS universal AP Reagent kit (ACD; 323250). Probes targeting *Ppib* (ACD; 313919) and *DapB* (ACD; 312039) mRNA were used as positive and negative controls, respectively. Staining of tissue and RNA detection was automated using a Discovery Ultra IHC/ISH staining platform (Roche Diagnostics). Briefly, slides were deparaffinized and treated with target retrieval buffers and a protease solution to free RNA from protein complexes before being incubated with target probes and sequential rounds of signal-amplifying oligonucleotides (all reagents provided by Advanced Cell Diagnostics).

Whole Slide Images (WSIs) of each section at 40x magnification were obtained using a NanoZoomer S360 (Hamamatsu Photonics K.K.). Manual annotation of frontal regions of each section, automated cell detection and quantification of punctae within detected cells were performed using QuPath v0.3.2^34^.

### Patients samples

Patients with a definite (i.e. neuropathologically confirmed) or probable diagnosis of sCJD according to World Health Organization criteria were recruited by the National Prion Clinic (London, UK). The study was approved by the London—Harrow Research Ethics Committee (reference 05/Q0505/113). Samples were obtained with written informed consent from controls, patients, or a patient’s consultee in accordance with applicable UK legislation and Codes of Practice. We selected 10 individuals from archived tissue collected by the National Prion Clinic and stored in our Unit. Our selection criteria included the final diagnosis, which was sporadic CJD, availability of frozen frontal cortex samples, storage of samples in histopathology cassettes that enable us to identify the different anatomical regions of the frontal cortex more easily, and codon 129 methionine homozygous *Prnp* genotype. For our control group, we included frontal cortex samples from individuals with low-level AD pathology or pathological ageing provided by the Queen Square Brain Bank. These samples (N = 10) were matched for sex (male = 4 sCJD and 5 controls; female = 6 sCJD and 5 controls) but not age (mean age for sCJD = 70.4 years, SD = 8.6; mean age for controls = 83.7 years, SD = 8.6) More information regarding the clinicopathological variables of the selected patients can be found in Supplementary Table 2.

We were also able to source 3 non-dominant frontal lobe biopsy samples from sCJD patients. These extremely rare samples have been collected over 20 years by the National Prion Clinic because the differential diagnosis of CJD sometimes requires excluding neuroinflammatory conditions like primary cerebral vasculitis. Occasionally, this can only be determined through histological examination of brain tissue in life. These samples offer two main advantages: they are well preserved, and there is no post-mortem delay since tissue archiving is fast, usually less than 30 minutes after sample collection. The control samples for this group included frontal lobe biopsies from non-neurodegenerative disease controls with mixed clinical diagnoses and only non-specific minor histological changes (pathological non-diagnostic samples), provided by BRAIN UK. These samples (N = 3) were sampled similarly to the biopsies and individuals were matched for sex (male = 2 sCJD and 2 controls; female = 1 sCJD and 1 control) while the age was matched only partially (mean age for sCJD = 56.6, SD = 13.3; mean age for controls = 60.6, SD = 3.3). More information regarding the clinicopathological variables can be found in Supplementary Table 2.

### DNA extraction from prion-infected frozen brain tissue

50-100 mg of brain tissue were transferred to a 2 mL screw-cap tube (Alpha Laboratories; CP5932). 450 μL ATL lysis buffer (QIAGEN; 939016) and 50 μL proteinase K 20 mg/mL (Invitrogen; AM2548) were added, and tubes were left in a Thermomixer Comfort heating block (Eppendorf) overnight at 50°C with mixing at 800 rpm. The next day, 500 μL of TRIS-equilibrated phenol (Sigma-Aldrich; P4557) were added and mixed by inversion. The tubes were centrifuged at 16,000 g for 5 min at room temperature before transferring the upper aqueous phase to a fresh tube and discarding the lower organic phase. The addition of phenol, centrifugation and keeping of the aqueous phase was repeated. 500 μL of a 1:1 mix of TRIS-equilibrated phenol and chloroform mixture were added and mixed by inversion. After centrifugation, the aqueous phase was transferred to a fresh tube and 500 μL chloroform was added. After centrifugation, the aqueous phase was transferred to a fresh tube and removed from BSL-3 facilities to BSL-2 facilities. 500 μL of 100% cold ethanol was added to induce DNA precipitation. The supernatant was then aspirated and discarded without disturbing the DNA pellet, which was left to dry for a couple of minutes and was then resuspended in water.

### *PRNP* codon 129 genotyping

TaqMan^®^ SNP Genotyping Assays with appropriate probes were used for *PRNP* codon 129 genotyping according to the manufacturer’s instructions. Briefly, 5 μL of TaqMan™ Genotyping Master Mix (Applied Biosystems; 4371353), 0.5 μL of assay probes (Thermo Fisher Scientific; 4351379; assay ID: C_2969398_10), 1 μL DNA and 3.5 μL water were added in each well of a MicroAmp™ Fast Optical 96-Well Reaction Plate (Applied Biosystems; 4346906). The plate was sealed, vortexed and spun down and then placed in a QuantStudio 12K Flex Real-Time PCR System (Thermo Fisher Scientific) where the following PCR program was run: 1 cycle of 10 min at 95°C, 40 cycles of 15 s at 95°C and 1 min at 60°C. End-point fluorescence was detected, and the allelic discrimination plots were used to identify the sample genotype. MM, MV, VV, and negative controls were included in each run.

### RNA extraction from prion-infected frozen brain tissue

Frozen frontal cortex brain slices were left to thaw and were hand-dissected. Approximately 50-100 mg of the superior frontal gyrus was homogenised in Tri Reagent (Zymo Research) using a pellet pestle (Kimble) and incubated at room temperature for 2 hours. Total RNA was purified using the Direct-zol RNA miniprep kit (R2050; Zymo Research) following the manufacturer’s instructions. DNAse I treatment was performed in tube followed by column purification (Zymo Research). All RNA samples were subsequently run on a TapeStation 2200 (Agilent, Santa Clara, US) using High Sensitivity RNA ScreenTapes.

### Total RNA libraries preparation and sequencing

100 ng of Total RNA was used as input material to pepare librairies using the Universal Plus Total RNA-Seq with Human AnyDeplete (Tecan) according to the manufacturer’s instructions. Resulting libraries were visualized and quantified on the 2200 TapeStation Instrument and diluted at 2nM concentration and pooled in two batches of 7 libraries each. Each pool was sequenced on the Illumina NextSeq 500 after the addition of 5% PhiX sequencing control library (FC-110-3002; Illumina) using the following settings: paired-end, read 1: 75 nt, read 2: 75 nt.

### Total RNA-sequencing data analysis

The fastq files were aligned to the human reference genome (GRCh38) using the STAR aligner and resulting alignments were converted to binary format and sorted using samtools. Gene features were counted using the GenomicAlignments library in R and the resulting SummarizedExperiment was loaded into DESeq2 using the design formula: ∼ condition. DESeq2 was used for differential expression testing using the default settings.

## Data availability

All sequencing data that support the findings of this study have been deposited in the National Center for Biotechnology Information Gene Expression Omnibus (GEO) and will be accessible through the GEO website.

## Author contribution

S.M. had full access to all data in the study and takes responsibility for integrity of the data and accuracy of the data analysis. Concept and design: E.A.V. and S.M.; acquisition, analysis or interpretation of data: A.D., F.Z., T.T., T.T., T.N., C.S. and E.A.V.; drafting of manuscript: A.D., E.A.V. and S.M.; obtaining funding: E.A.V. and S.M.; study supervision: S.M. and E.A.V.

