## Extended Data for "Single-nuclei transcriptomics of mammalian prion diseases identifies dynamic gene signatures shared between species"

### Extended Data Figure 1

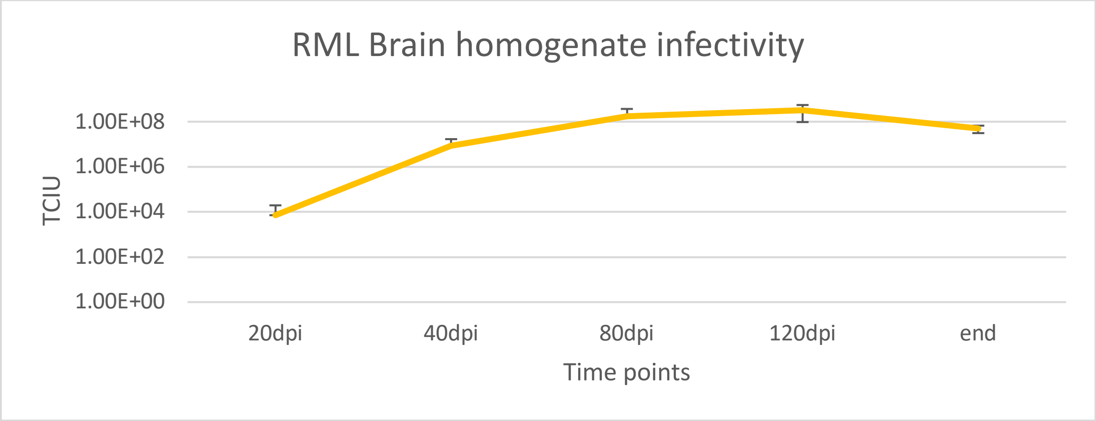


***Extended Data Figure 1: The scrapie cell assay validates the effective selection of the 5 time points for the mouse transcriptomics study.*** *The mean of infectious units from the same samples (n = 3) is plotted on a logarithmic y-axis. Error bars indicate the standard deviation. TCIU: tissue culture infectious units, referred to as infectious units in the text.*

### Extended Data Figure 2

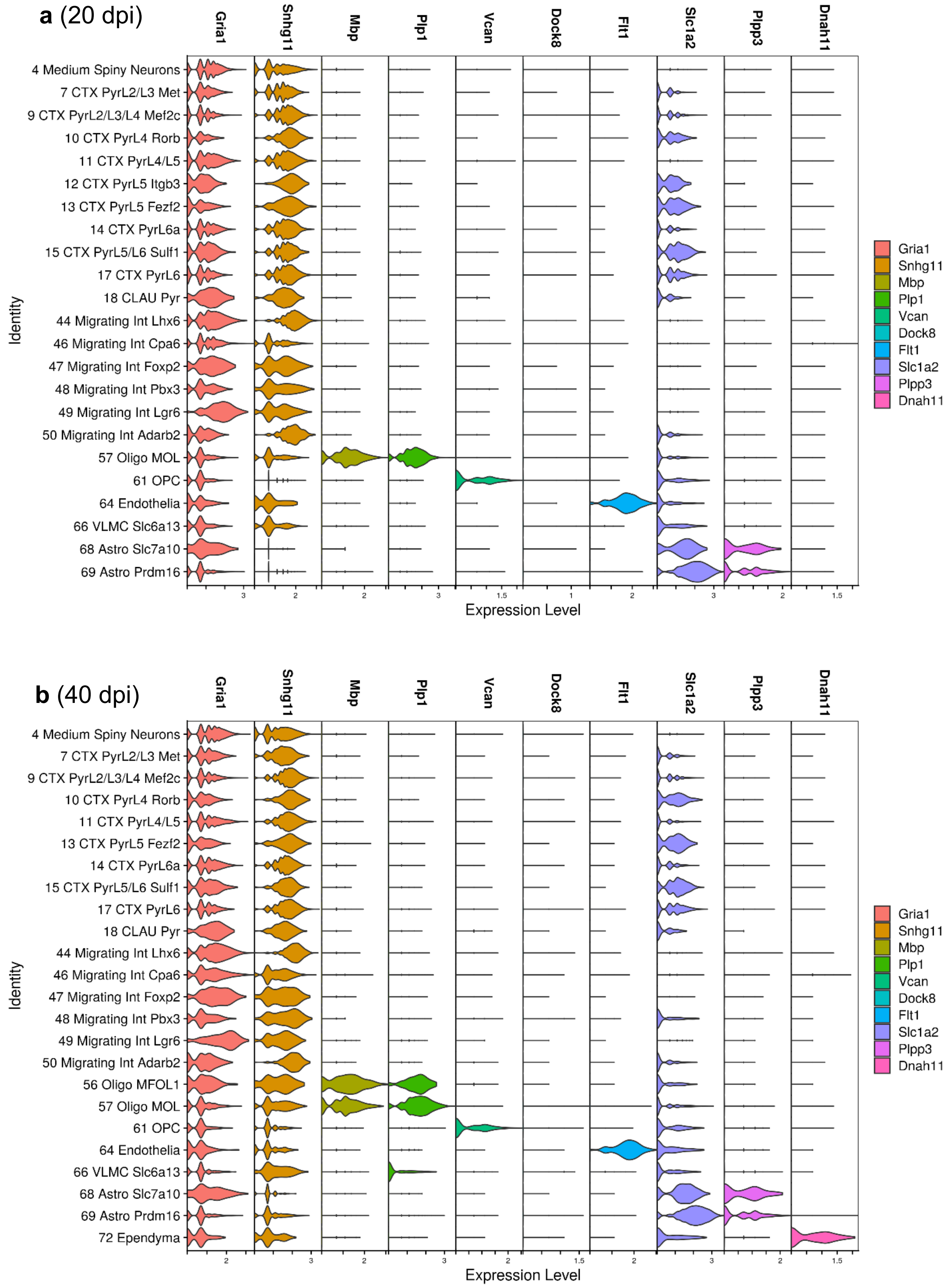

**c** (80 dpi)

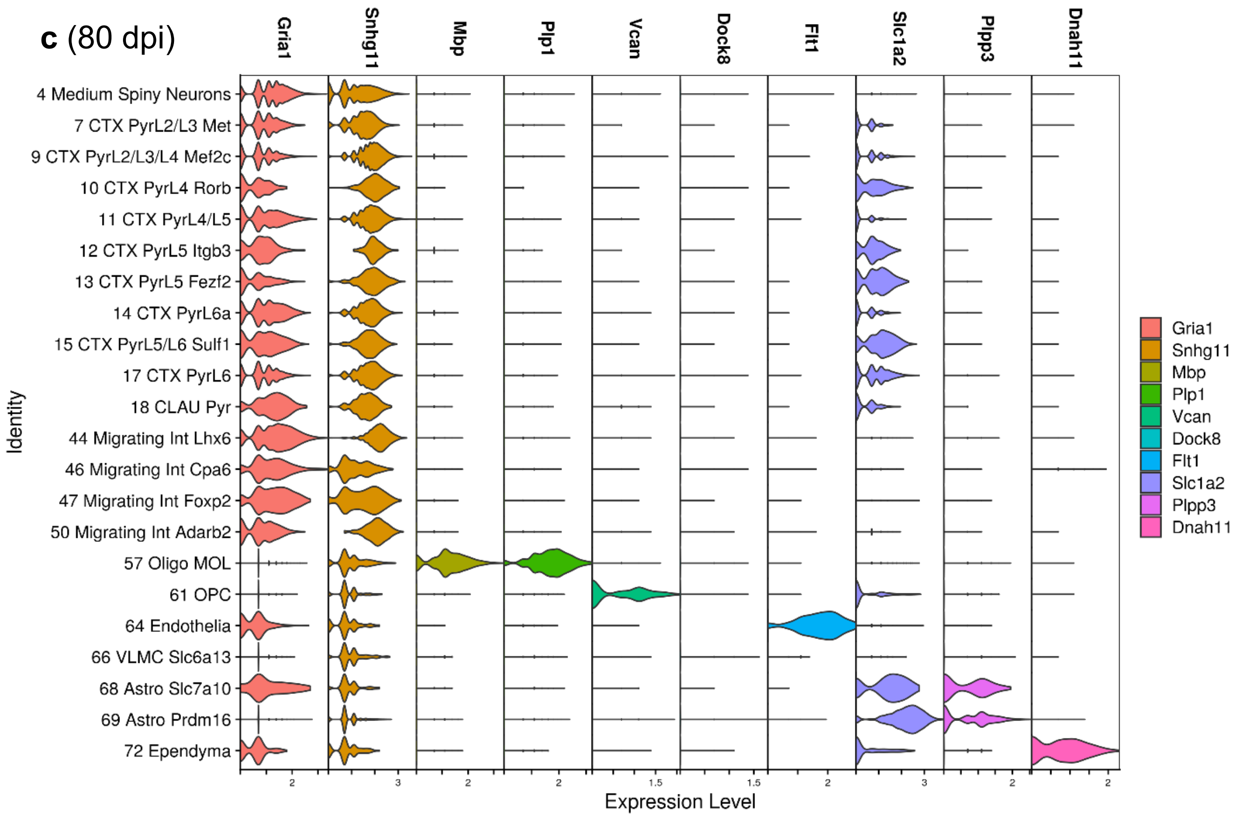


**d** (120 dpi)

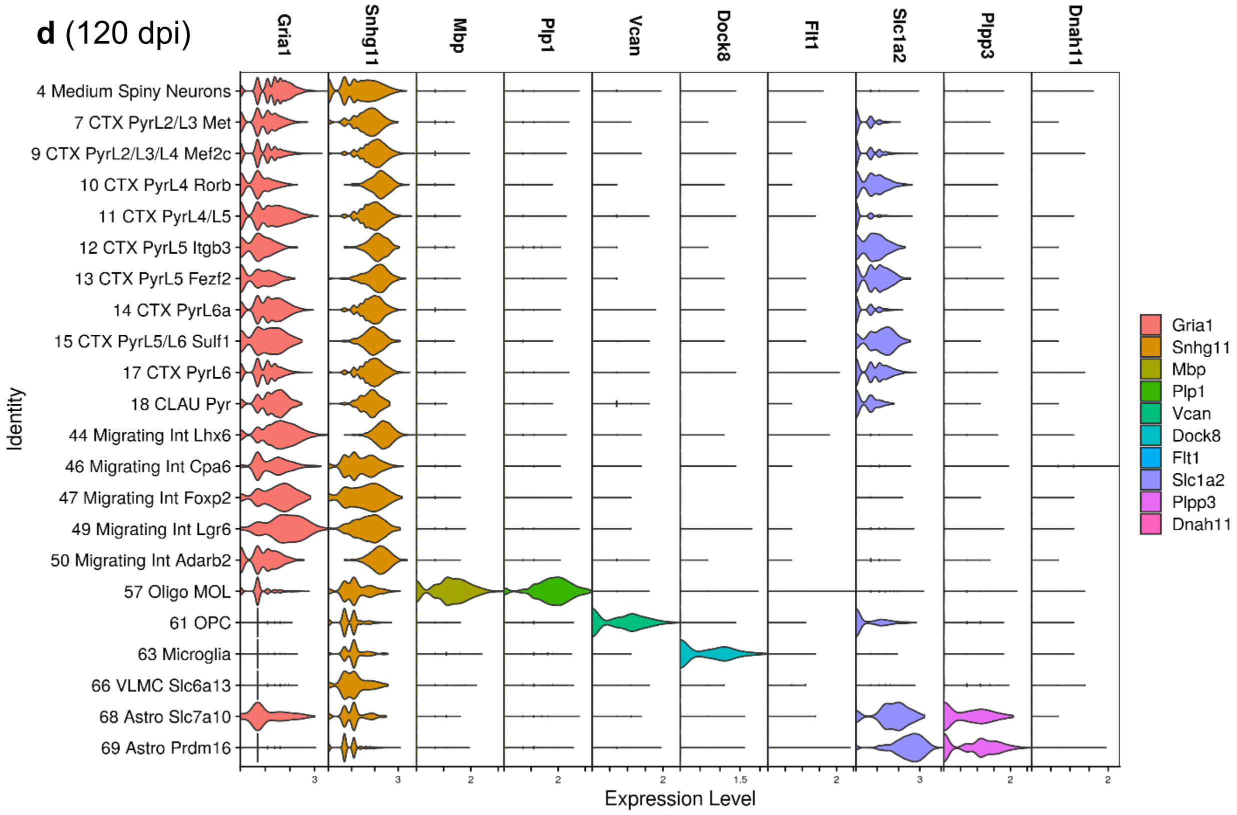

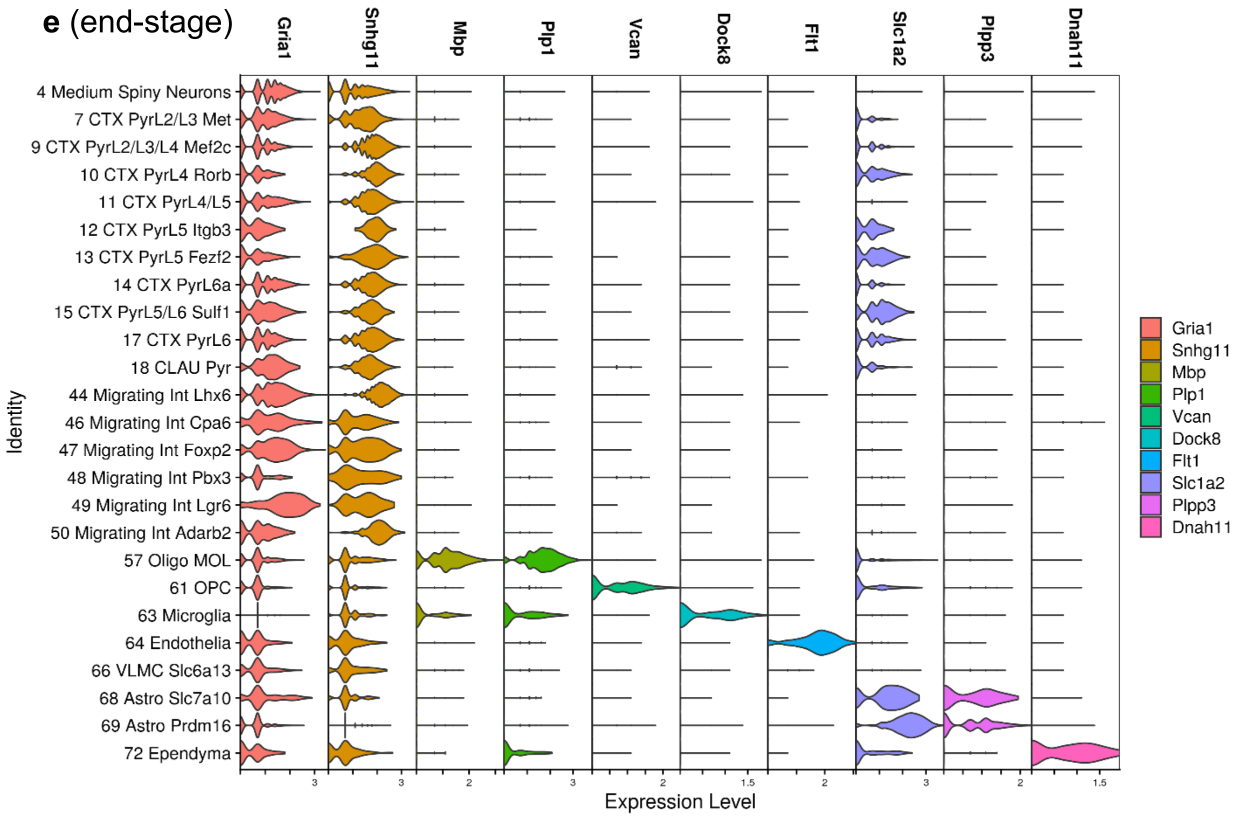


***Extended Data Figure 2: The expression of known marker genes corroborates cluster identities.*** *Violin plots of the expression of known marker genes across the identified clusters in 5 time points. Marker genes used: Gria1 and Snhg11 for neurons; Mbp and Plp1 for oligodendrocytes; Vcan for oligodendrocyte precursor cells; Dock8 for microglia; Flt1 for endothelia; Slc1a2 and Plpp3 for astrocytes; Dnah11 for Ependyma. The expression level corresponds to the sctransform normalised expression values of the dataset.*
